# STUDIES ON INTERACTION OF MORIN WITH CURLI TO IDENTIFY AGGREGATION NATURE

**DOI:** 10.1101/2025.03.25.645188

**Authors:** Tirthankar Roy, Nisha Bhat, Shritoma Sengupta, Samudra Prosad Banik, Maitree Bhattacharyya, Pijush Basak

## Abstract

Morin (3,5,7,2′,4′-pentahydroxyflavone) is a flavonoid compound exhibited a range of biological functions, including anti-inflammatory, antioxidant, antiviral, and anti-allergic activities. It has the ability to disaggregate amyloid fibrils, making it a potential inhibitor of diseases caused by amyloid protein fibrillation. This study focuses on the interaction between Morin and Curli amyloid fibrils, which are produced by some Enterobacteriaceae, including *Salmonella* sp. And *E. coli*, as a major component of their biofilms. The Curli amyloid fibrils can lead to neurodegenerative diseases through a signalling cascade causing various human diseases, such as Alzheimer’s and Huntington’s diseases. Thioflavin T (ThT) fluorescence assays to purified Curli, and molecular docking between Morin and Curli indicates nature of binding at molecular level. The results showed that Morin quenched Curli fluorescence in a concentration-dependent manner, indicating dynamic quenching. Stern-Volmer analysis revealed that Morin binds to Curli with a binding constant of approximately 10^2^ L/mol. This binding was found to be spontaneous, with hydrophobic interactions playing a significant role. ThT fluorescence assays confirmed that Morin inhibited Curli fibrillation, suggesting a potential therapeutic application. Molecular docking analysis revealed specific interactions between Morin and Curli, providing insights into their binding at the molecular level. In conclusion, this study demonstrates the dynamic quenching of Curli fluorescence by Morin and suggested that Morin may inhibit Curli fibrillation, making it a valuable candidate for the development of therapeutic drugs.

## 1. Introduction

Morin (3,5,7,2′,4′-pentahydroxyflavone) is a flavonoid compound which generally found in osage orange, guava, tea and wine [1]. It exhibits biological functions such as anti-inflammatory activity antioxidant activity, antiviral activity ant allergic activity [2]. Their ability to disaggregate amyloid fibrils [3] helps to inhibit diseases that causes by the fibrillation of amyloid proteins. Morin also considered as an anti-cancerous agent for their action.

Some *Enterobacteriaceae* like *Salmonella sp* and *E. coli* produce Curli amyloid fibrils as a major component of their biofilm for their microbial lifestyle. This Curli protein has two different operons in their genes csgBAC and csgDEFG which synthesize seven proteins: csgA, csgB, csgC, csgD, csgE, csgF, csgG in which csgA and csgB are major proteins. This amyloid fibril formation is caused by aggregation of Curli protein which causes neurodegenerative disease via signalling cascade. The formation of amyloid fibrils causes several diseases in humans, including Alzheimer’s disease, Huntington’s disease, and prion diseases, although the process of amyloid formation in vivo is not well understood [4,5].

In this study it was analysed that Morin had some preservative properties for some fruits like banana. PCA analysis indicated that a significant effect of morin on metabolite changes occurred after 10 days storage, including the delayed accumulation of sucrose, fructose, α-d-glucose and β-d-glucose, the accelerated accumulation of alanine and GABA, the retarded formation of valine and l-aspartic acid, the suppressed degradation of saponin a. It suggested that morin is a promising preservative for banana to maintain the quality and extend shelf life. Also, this help morin to delay the skin colour of banana and change from green to yellow [6].

## 2. Materials and Methods

### 2.1. Chemicals

Thioflavin T (ThT) and YESCA agar was purchased from HIMEDIA, India and Buffer and chemicals were made by the ingredients purchased from HIMEDIA, India also. Fluorescence measurements were carried out using a Fluorescence spectrophotometer (Hitachi F-7100). Experiments were performed in 50mM 1N NaCl PBS buffer (pH 7.0) unless mentioned specifically. Morin were bought from Merck, India. Other components Kanamycin, Lysosyme, RNase A, DNase 1, MgCl_2_, Tris, SDS were from Sigma, India.

### 2.1. Purification of Curli

*E. coli* PHL628 cells are grown in YESCA broth with Kanamycin at 26°C and under 200rpm shaking for 48hrs. Bacterial pellets were collected by centrifugation and suspended in 10mM Tris-HCl at pH 8.0. After that the suspension was treated with a mixture of 0.1mg/ml RNase A, 0.1mg/ml DNase, and 1mM MgCl_2_ for 20mins at 37°C. Bacterial cells were then broken by sonication (30% amplification for 30s twice). After that Lysozymes were added (1mg/ml) and samples were incubated at 37°C. After 40 min, 1% SDS was added, and the samples were incubated for 20 min at 37°C with shaking (200 rpm). After this incubation, Curli was pelleted by centrifugation (10,000 rpm for 10 min at 8°C) and then suspended in 10 ml Tris-HCI (pH 8.0) and boil for 10 min. A second round of enzyme digestion was then performed as described above. Curli was then pelleted, washed in Tris-HCI at pH 8.0, and were suspended in 2X SDS-PAGE buffer and boil for 10 min. The samples were then electrophoresed on a 12% separating to 5% stacking gel run for 5h at 20 mA (or overnight at 100 V). Fibrillar aggregates were too large to pass into the gel and therefore remain within the well of the gel and could be collected. Once collected, the curli aggregates were washed three times with sterile water and then extracted by being washed twice with 95% ethanol. Purified curli were then resuspended in sterile water. Curli concentration was determined by Bradford reagent [7].

### 2.2. Binding with Curli at different concentration of Morin

The concentration of Curli was fixed at 7×10^-5^M and Morin concentration was varied from 0 to 4.5×10^-5^M at increments of 0.5×10^-5^M. The emission was checked at excitation wavelength of 295nm (Tryptophan) and emission wavelength was observed from 300nm to 500nm. Morin was prepared in PBS buffer. Temperature was maintained at 300K while mixing the both solutions [2].

### 2.3. Thioflavin-T (ThT) fluorescence assay

Curli (10×10^-5^M) was incubated for fibrillation in the presence and absence of Morin (10×10^-5^M). 5×10^-5^M ThT was added to both incubated solutions. Emission spectra (460-550nm) of these two was recorded using excitation at 450nm [8].

### 2.4. Docking

Three-dimensional structure of Curli was collected from Protein Data Bank (PDB) and Three-dimensional structure of morin collected from PubChem. Morin was docked through CB Dock. It predicted specific protein binding sites and calculated centers and sizes with a new curve-based cavity detection method and performed docking using the popular docking program Autodock Vina. This method was carefully optimized and achieved a ∼70% success rate [9]. After getting the docking result in .pdb format, its further uploaded in Protein-Ligand Interaction Profiler (PLIP web tool) for more specific data to visualizes and detected how they interaction each other’s, their bond angles and their bond distance [10]. Finally, the conformation was visualized in PyMol.

## 3. Results

### 3.1. Effect of Morin on Curli fluorescence characteristics

Fluorescence quenching was differentiated as dynamic and static quenching. They both differed from each other by their dependency on temperature and viscosity. In this experiment Curli solution were stabilized at 7×10^-5^M and the concentration of morin varied from 0 to 4.5×10^-5^M. The effect of Morin on Curli fluorescence intensity at 300K was shown in Fig.1. It was concluded that a continuous decrease in fluorescence intensity was a result of fluorescence quenching. Also, Morin did not show any fluorescence emission at 295nm excitation. The phenomenon of fluorescence quenching was stated by well-known Stern-Volmer equation (2)

**Fig1:**
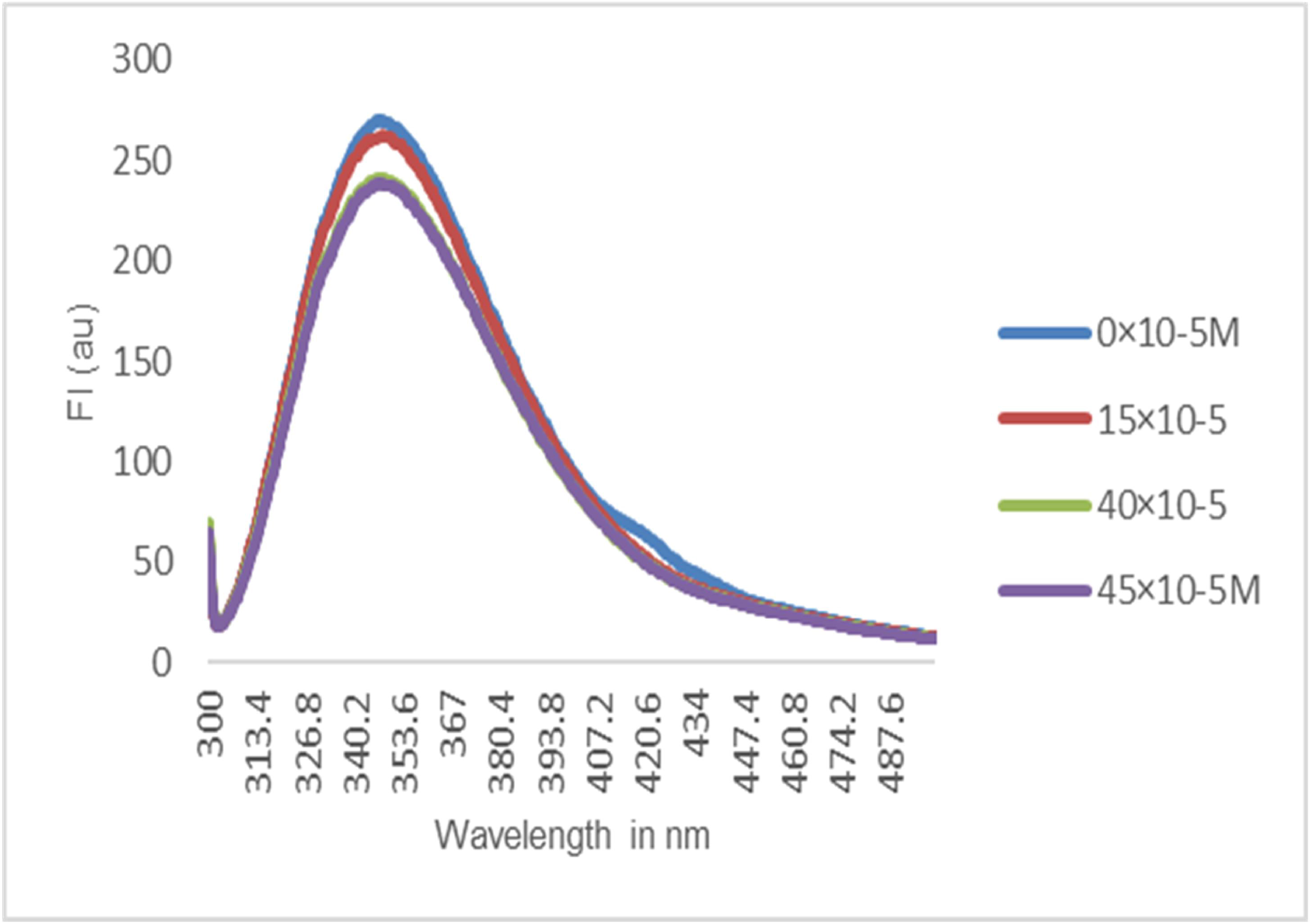
Emission spectra of Curli in the presence of various concentrations of Morin. (T= 300K, _□ ex_ = 295nm)

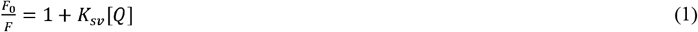

In above equation F_0_ and F were fluorescence intensity in the absence and presence of quencher [Q]. K_sv_ was the Stern-Volmer quenching constant and [Q] was concentration of quencher. Stern-Volmer equation was implied in Fig 2. and the data were compiled in Table 1 to estimate Stern-Volmer quenching constant. The result showed that fluorescence quenching by Morin at different concentration. Observation shows K_*sν*_ was directly proportional to quencher’s concentration and graph (Fig 2) showed a curve nature of K_*sν*_ which indicated that the both static and dynamic quenching occurred between Curli and Morin.

**Table 1:**
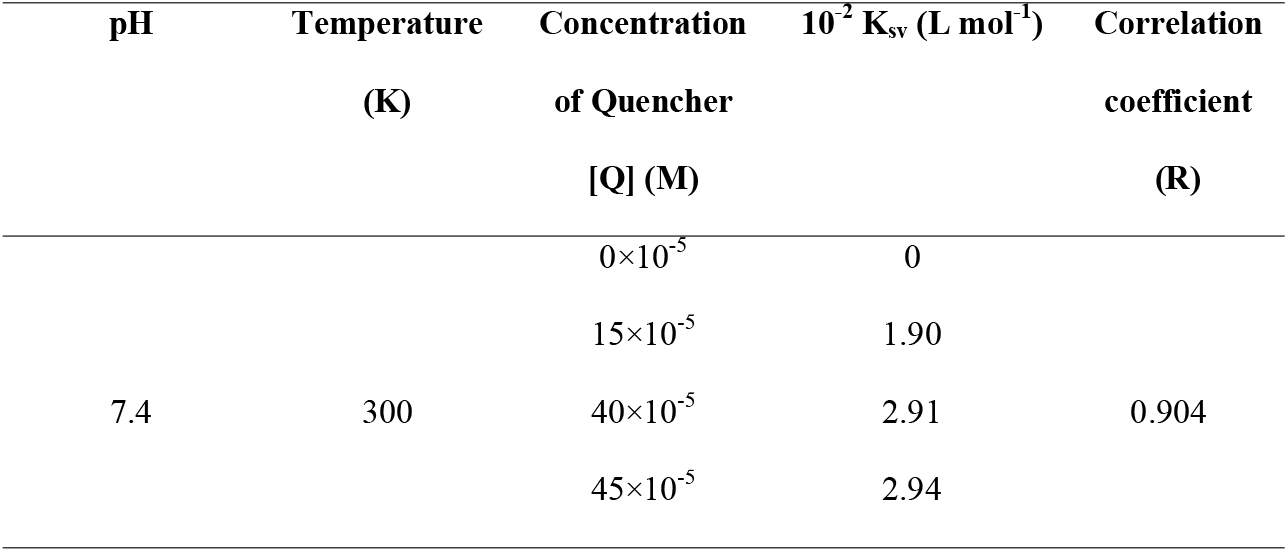
Stern-Volmer quenching constant for the interaction of Morin with Curli at various concentration.

**Fig2:**
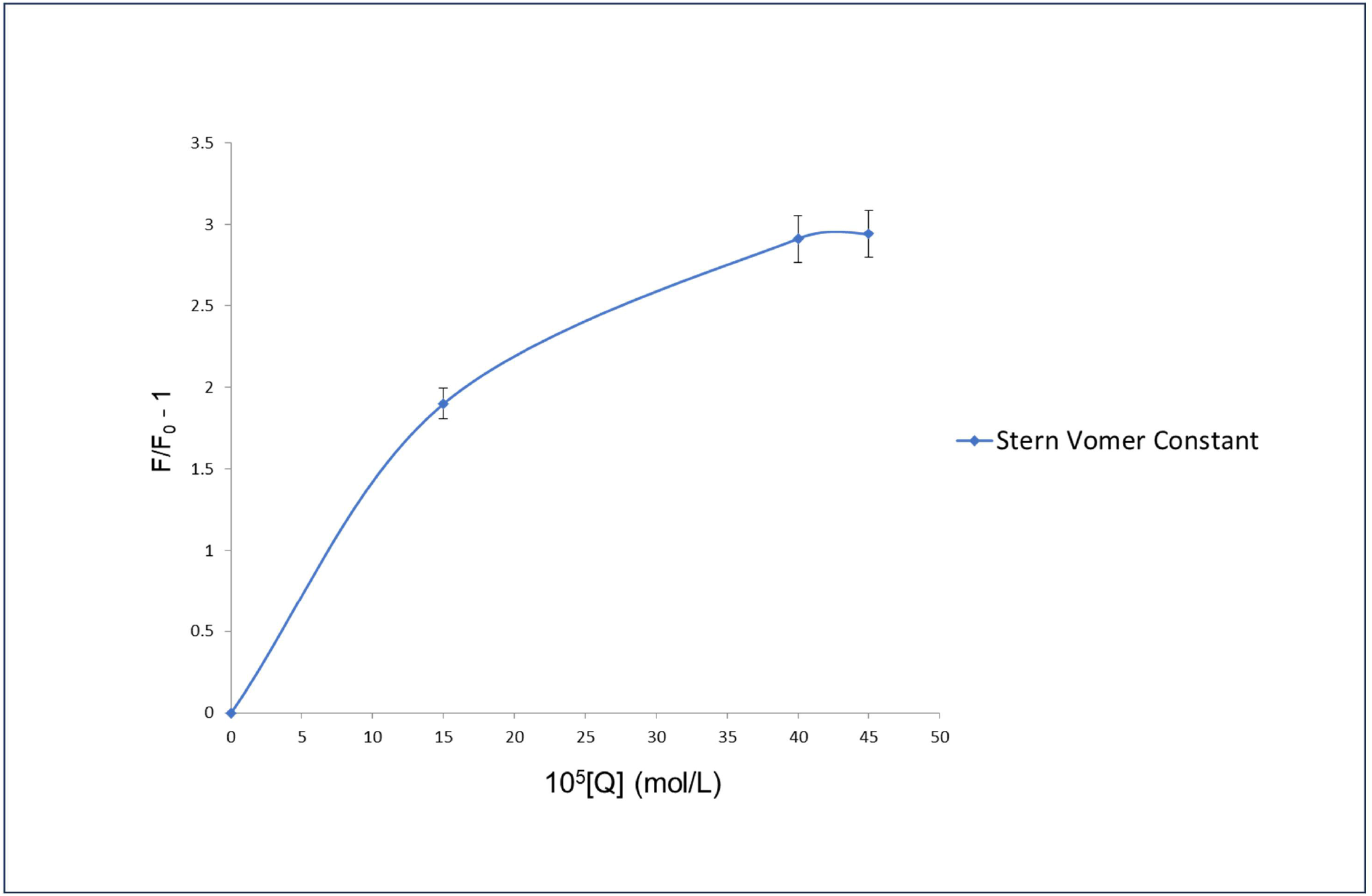
Plots (F_0_/F – 1) vs [Q] for elucidation of the quenching constant K_sv_ for Morin-Curli binding interaction from fluorescence data. Values are the average of results from triplicates trials, error bars indicate the SD values.

### 3.2. Binding parameters between Morin and Curli

To seek the quantification of the observed quenching of fluorescence of Morin with Curli modified Stern-Volmer equation was used instead of conventional form of the equation considering the complex structure of Curli. The modified stern volmer equation was [11]:

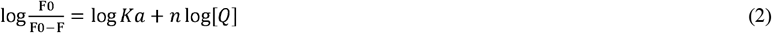

In the above equation Ka was the effective quenching constant and n was number of binding sites. From this Ka value was obtained and the value was directly used to determine the free energy change using the equation [12]:

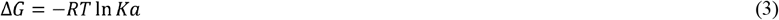

Fig 3 showed plots for Morin-Curli system at different concentration. The corresponding results were concluded in Table 2. The increasing trend of Ka with increasing concentrations were in accordance with Ksv ‘s dependence on concentration was mentioned in “Binding with Curli at different concentration of Morin”

**Table 2:**
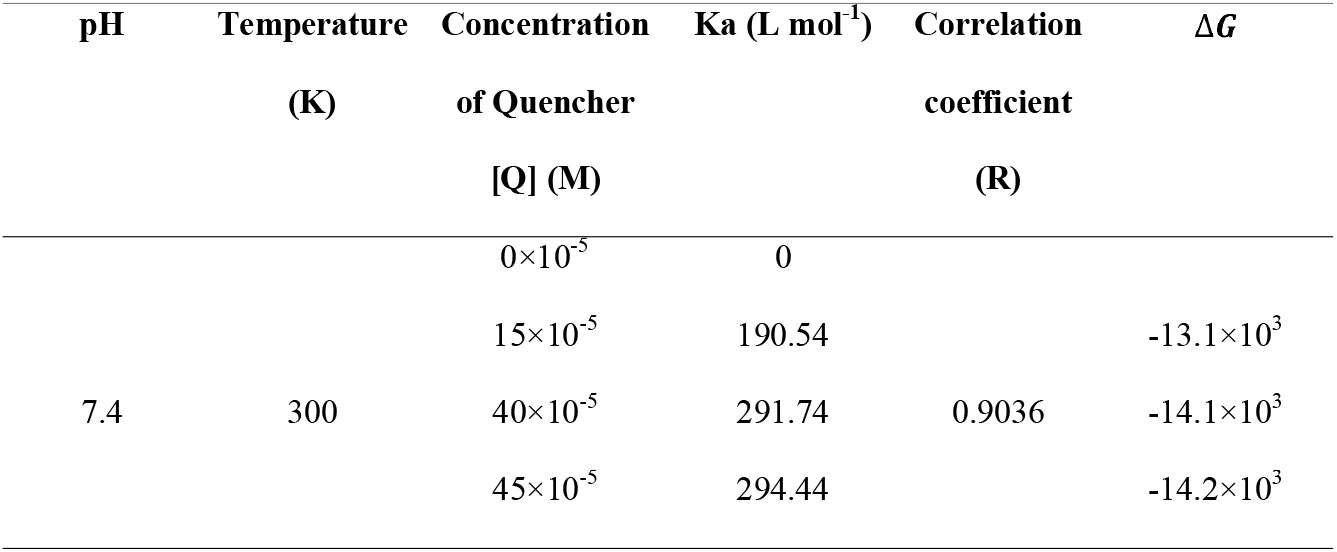
Modified Stern volmer quenching constant for the interaction of Morin with Curli at various concentration. R is the correlation coefficient

**Fig3:**
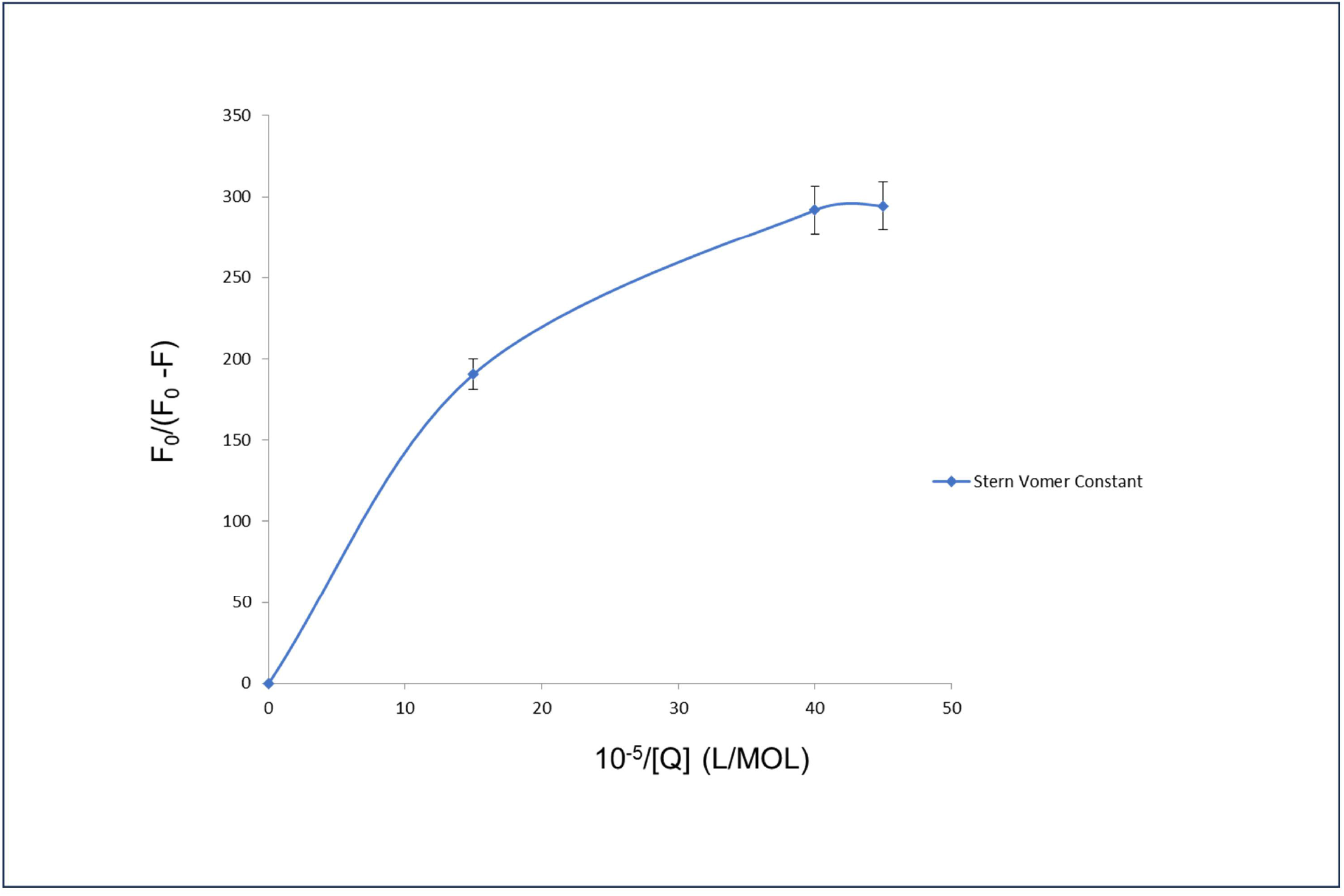
Modified Stern-Volmer plot. Values are the average of results from triplicates trials, error bars indicate the SD values

### 3.3. Thioflavin T (ThT) fluorescence

ThT was used as a probe to detect amyloid formation [13]. The mechanism was depended on the increasing fluorescence intensity when ThT binds to the amyloid fibrils. In this experiment emission intensity increased 2.44-fold when ThT added with Curli in PBS (pH 7.4) at 37°C for 100hrs under shaking conditions (160-180rpm). But for Native Curli, emission did not change. To explore the inhibitory effect of Morin against Curli was done in absence and presence of Morin. In Fig 4a the ThT emission intensity (490nm) was increasing in absence of Morin as Curli forms fibrils. But in presence of Morin with the Curli, slight inhibition of fibrillation was noticed in respect of time (Fig 4b). Also, when Morin concentration was increased, there was decrease of ThT fluorescence intensity (Fig 4c)

**Fig4a:**
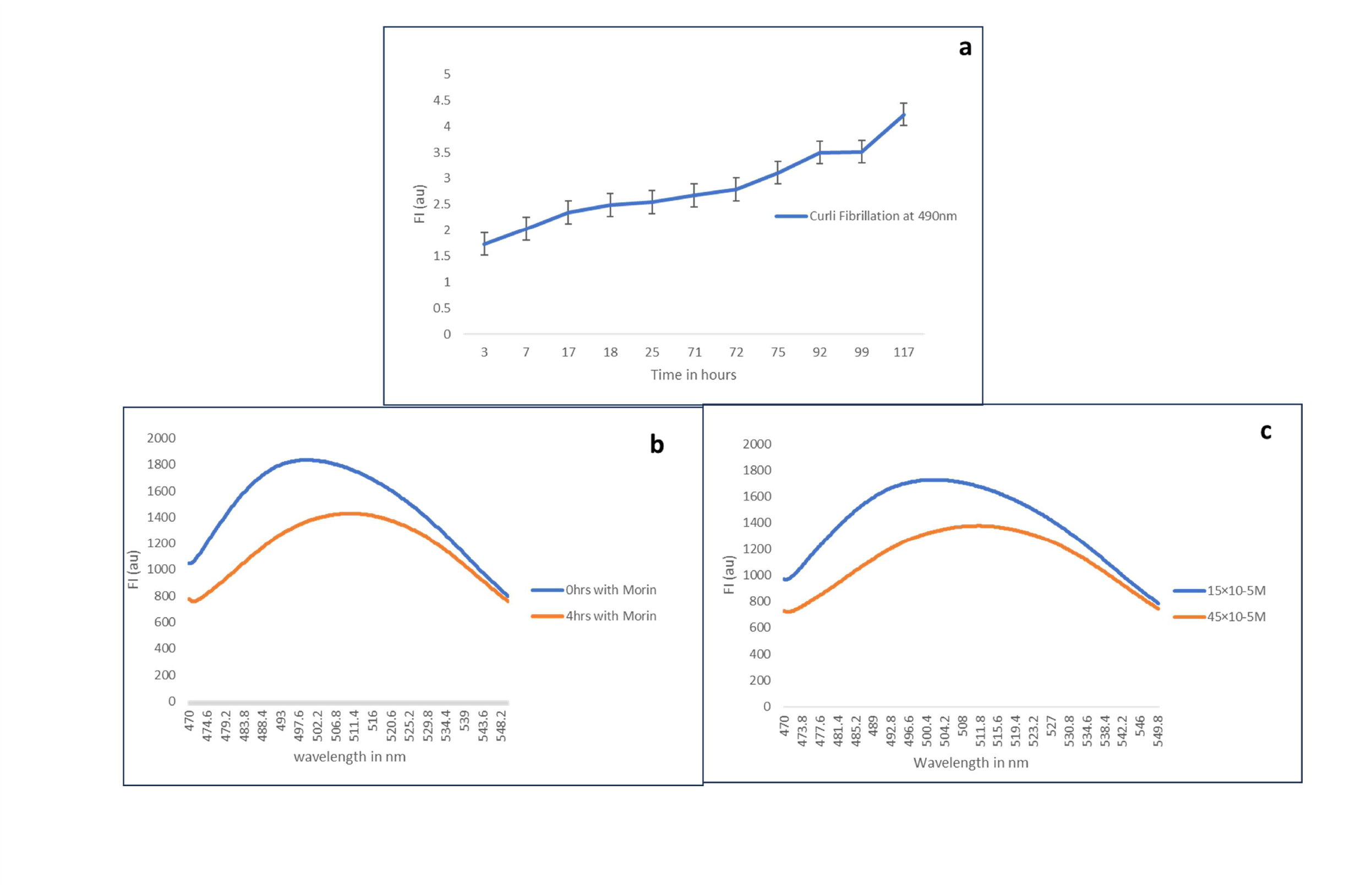
Curli fibrillation assay by binding with **(a)** Thioflavin T, **(b)** Thioflavin T in presence of Morin and **(c)** Thioflavin T in presence of different concentration of Morin. (T=300k, Excitation wavelength: 450nm; Emission wavelength 470nm-550nm).

**Fig5:**
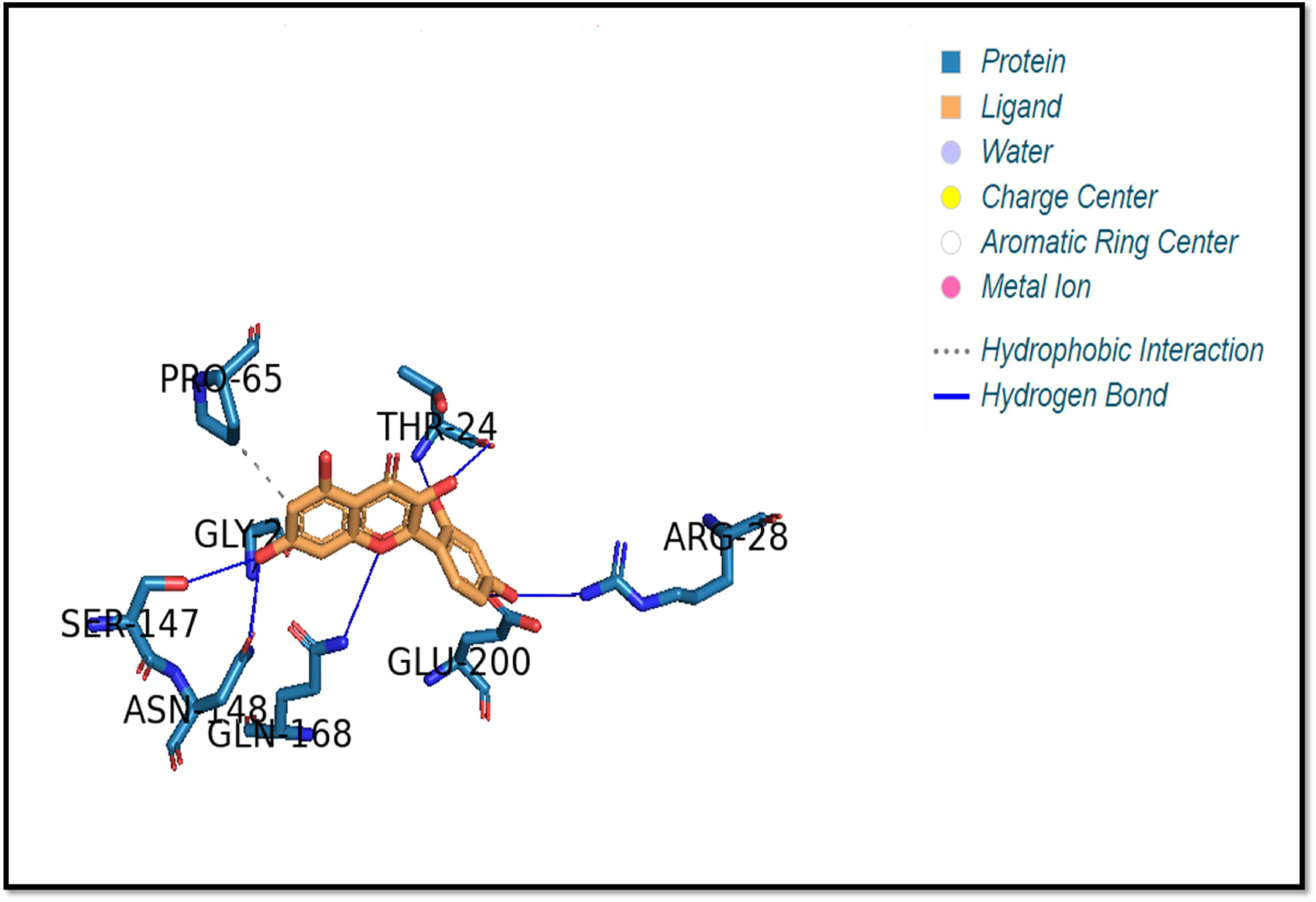
Docked conformation of Morin with Curli obtained from CB Dock. Interacting residues of protein are also shown.

### 3.4. Docking

For corroborating the observations, Curli was docked with morin and energetically best bound conformation of Morin in the protein structure was shown in Fig5. The stability of this protein-ligand conformation was thought to be depend upon their inter and intra-domain interaction. It was clear to say that, from Fig4. the polar atoms of some residues from both domains interact with oxygen atoms of Morin. The amino acid residues of Curli protein GLY 21, THR 24, ARG 28, SER 147, ASN 148, GLN 168 and GLU 200 mainly favour the binding with Morin. Moreover, these residues were in hydrogen bonding with Morin (Table 3). From the docked structured we have also found that residues PRO 65 which exhibits hydrophobic interaction with Morin.

**Table 3:**
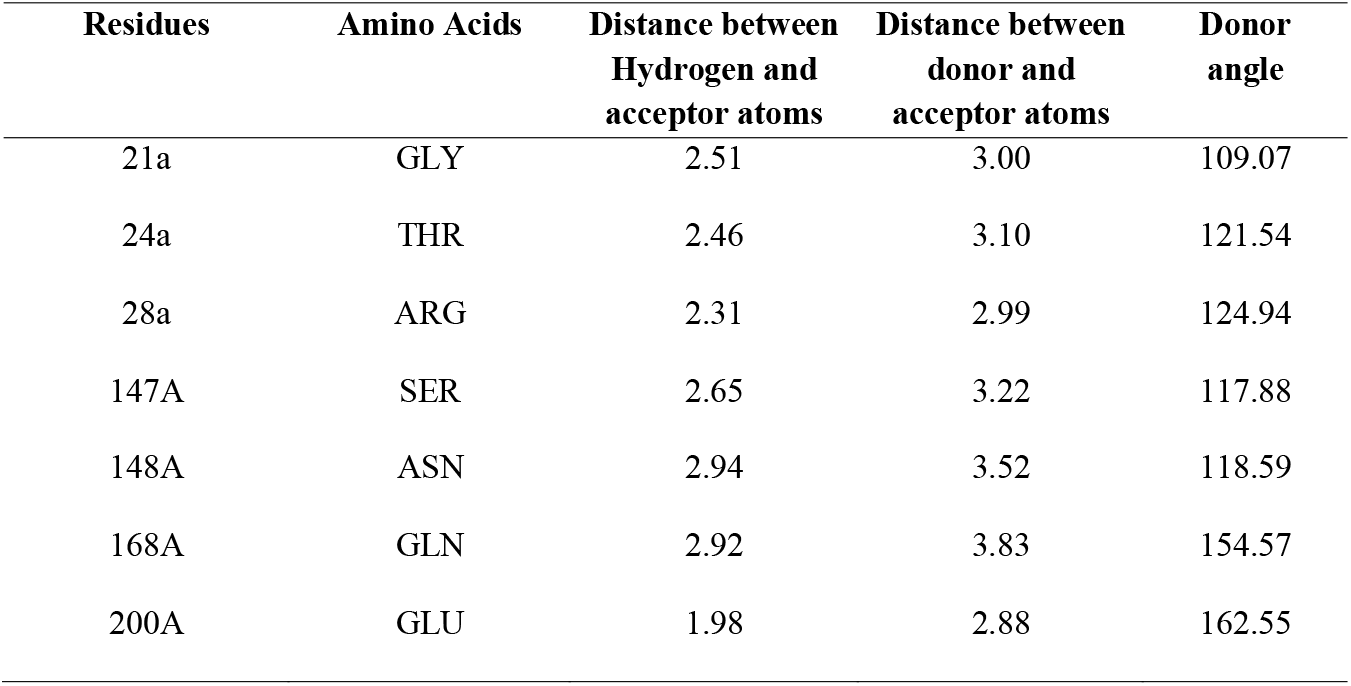
Distance and donor angle between the interacting residues of Curli and Morin as btained from PLIP webtool.

## 4. Discussion

Experimental data suggested that both static and dynamic quenching occurs during the binding of Curli with Morin. Static quenching caused due to formation of ground-state complex between protein fluorophores and Morin. The dynamic quenching, also known as collisional quenching, resulted from collision between protein fluorophores and Morin [14,15]. Also, both static and dynamic quenching concluded that binding between Curli and Morin favourable in both ground state and entire excited state. The ground-state complex could be reasonably stable, for frequently observed both static and dynamic quenching. Dynamic quenching also decreased the mean decay time of the entire excited-state [16].

Observed value of modified Stern-Volmer constant (Ka) in Gibbs free energy change equation (in Table 2), it was interestingly found out that the free energy change is negative indicating favourable interaction between Curli and Morin and also indicating that binding to Morin is highly contributed by positive entropy change. High *ΔG* value also revealed that the binding is very facile from thermodynamic aspect which again gave the idea of stable interaction between Curli and Morin [12].

During Thioflavin-T (ThT) assay, the ThT bound to amyloid fibrils gave off a strong fluorescence signal which helped to quantify the formation of amyloid fibrils [13,17,18]. The Morin bound to Curli residues had greater aggregation propensity and prevented structural conformational changes that led to amyloid fibrillation. Morin bound to the fibrillar conformation of Curli by hydrogen bonding and hydrophobic interaction throughout the protein surface and stabilized them. It also inhibited the release of oligomeric species which could be nuclei or template for further fibrillation [19]. The docking results also confirmed that the micro-environment of the interaction between Curli and Morin are thermodynamically favourable and prefer weak interactions.

## 5. Conclusion

The interaction between Curli and Morin was observed by fluorescence spectroscopy and docking software to identify the nature and location of interaction between Morin and Curli protein to explore the therapeutic application for potential anti-aggregative agent. The results indicated the quenching mechanism of fluorescence of Curli by Morin was a dynamic quenching. The binding parameters calculated from Stern-Volmer method showed that morin binds to Curli with the binding affinity of the order of 10^2^ Lmol^-1^. The binding was spontaneous and hydrophobic interaction played a major role. This study also showed Morin can suppress the aggregation or fibrillation of the Curli protein resulting inhibition of protein aggregation mediated disorder. This interaction and inhibitory effect of Morin on Curli could be valuable for designing potential therapeutic drugs for future prospect.

## 8. Credit authorship contribution

**Tirthankar Ray**: Experiment, Validation, Writing - Original Draft, **Nisha Bhat:** Experiment, Writing Review & Editing. **Pijush Basak:** Supervision, Conceptualization, Resources, Validation, Formal and Data analysis, Writing - Review & Editing, **Maitree Bhattacharyya:** Project administration, Resources, Idea, **Shritoma Sengupta**: Writing - Review & Editing, **Samudra Prosad Banik**: Resources, Validation, Formal and Data analysis.

## 9. Data availability

Data will be made available on request.

## 10. Acknowledgement

Curli extracted from *E*.*coli* PHL628 which was kind donation from the laboratory of Prof. Anthony Hay, Cornell University, USA. The entire experiment was supported and conducted at Department of Biotechnology (Rosalind Franklin Biotechnology Laboratory), JBNSTS, Kolkata, West Bengal, India. Authors also like to thank Department of Science and Technology and Biotechnology, Government of West Bengal, India for the chemicals and the laboratory & instrumental facilities for entire experiment. Authors acknowledged Department of Microbiology, Maulana Azad College for their support.

## References

Hsiang, C. Wu, S. Ho, T. (2005) Morin inhibits 12-O-tetradecanoylphorbol-13 -acetate-induced hepatocellular transformation via activator protein 1 signaling pathway and cell cycle progression, Biochemical Pharmacology Volume 69, Issue 11, 10.1016/j.bcp.2005.03.008

Hu, Y. Yue, H. Li, X. Zhang, S. Tang, E. Zhang, L. (2012) Molecular spectroscopic studies on the interaction of morin with bovine serum albumin, Journal of Photochemistry and Photobiology B: Biology, Volume 112, Pages 16–22

Noor, H. Cao, P. Raleigh, DP. (2023) Morin hydrate inhibits amyloid formation by islet amyloid polypeptide and disaggregates amyloid fibers. Protein Sci. 2012 Mar;21(3):373–82. doi: 10.1002/pro.2023. Epub 2012 Jan 31. PMID: 22238175; PMCID: PMC3375438.

Barnhart, M. Chapman, M. (2006) Curli biogenesis and function. Annu Rev Microbiol.;60:131–47. doi: 10.1146/annurev.micro.60.080805.142106. PMID: 16704339; PMCID: PMC2838481.

Yan, Z. Yin, M. Chen, J. (2020) Assembly and substrate recognition of curli biogenesis system. Nat Commun 11, 241. 10.1038/s41467-019-14145-7

Zhu, H., Yang, J., Jiang, Y., Zeng, J., Zhou, X., Hua, Y., Yang, B. (2018) Morin as a Preservative for Delaying Senescence of Banana. Biomolecules. Jul 12;8(3):52. doi: 10.3390/biom8030052. PMID: 30002341; PMCID: PMC6164001.

Sivaranjani, M., Hansen, E. G., Perera, S. R., Flores, P. A., Tukel, C. and White, A. P. (2022). Purification of the Bacterial Amyloid “Curli” from Salmonella enterica Serovar Typhimurium and Detection of Curli from Infected Host Tissues. Bio-protocol 12(10): e4419. DOI: 10.21769/BioProtoc.4419.

Rana, S., and Ghosh, K. (2020): Inhibition of fibrillation of human γD-crystallin by a flavonoid morin, Journal of Biomolecular Structure and Dynamics. 10.1080/07391102.2020.1775701

Liu, Y., Grimm, M., Dai, Wt. (2020) CB-Dock: a web server for cavity detection-guided protein–ligand blind docking. Acta Pharmacol Sin 41, 138–144. 10.1038/s41401-019-0228-6

Adasme, M. and others, (2021) PLIP 2021: expanding the scope of the protein–ligand interaction profiler to DNA and RNA, Nucleic Acids Research, Volume 49, Issue W1, 2, Pages W530–W534, 10.1093/nar/gkab294

Paul, B.K. and Gucchait, N. (2011) Spectral deciphering of the interaction of an intermolecular charge transfer fluorescence probe with cationic protein: thermodynamic analysis of the binding phenomenon combined with blind docking study, Photochem. Photobiol. Sci. 10 980–991.

Basak, P., Debnath, T., Banerjee, R., Bhattacharyya, M. (2016). Selective binding of divalent cations toward heme proteins.

Biancalana, M., & Koide, S. (2010) Molecular mechanism of Thioflavin-T binding to amyloid fibrils. Biochimica et Biophysica Acta, 1804, 1405–1412. DOI: 10.1016/j.bbapap.2010.04001

Silva D., Cortez, C. Bastos J.K., Louro, S., (2004) Methyl parathion interaction with human and bovine serum albumin, Toxicology Letters, Volume 147, Issue 1, Pages 53–61, ISSN 0378-4274, 10.1016/j.toxlet.2003.10.014.

Bhattacharyya, M., Chaudhuri U., Poddar, R.K., (1990) Evidence for cooperative binding of chlorpromazine with hemoglobin: Equilibrium dialysis, fluorescence quenching and oxygen release study, Biochemical and Biophysical Research Communications, Volume 167, Issue 3, Pages 1146–1153, ISSN 0006-291X, 10.1016/0006-291X(90)90643-2.

Joseph R. Lakowicz, Principles of Fluorescence Spectroscopy, Third Edition, Chapter 8, ISBN-10: 0-387-31278-1 ISBN-13: 978-0387-31278-1.

Xue C., Lin, T., Chang, D., Guo, Z. (2017) Thioflavin T as an amyloid dye: fibril quantification, optimal concentration and effect on aggregation, Royal Society Open Science, Volume 4, pages 160696, 10.1098/rsos.160696

Hudson, S., Ecroyd, H., Kee, Tak W.; Carver, J. A., The thioflavin T (2009) fluorescence assay for amyloid fibril detection can be biased by the presence of exogenous compounds, 5960–5972. https://ro.uow.edu.au/scipapers/945

Patel, P., Parmar, K., Das, M., (2018) Inhibition of insulin amyloid fibrillation by Morin hydrate, International Journal of Biological Macromolecules, Volume 108, Pages 225–239, ISSN 0141-8130, 10.1016/j.ijbiomac.2017.11.168.

